# Structural basis of inhibition of a putative drug efflux transporter NorC, through a single-domain camelid antibody

**DOI:** 10.1101/2020.10.21.349639

**Authors:** Sushant Kumar, Arunabh Athreya, Ashutosh Gulati, Rahul Mony Nair, Aravind Penmatsa

## Abstract

Multi-drug efflux is a major mechanism of acquiring antimicrobial resistance among superbugs. In this study, we report the X-ray structure of NorC, a 14 transmembrane major facilitator superfamily member that is implicated in fluoroquinolone resistance in drug-resistant *Staphylococcus aureus* strains, at a resolution of 3.6 Å. The NorC structure was determined in complex with a single-domain camelid antibody that interacts at the extracellular face of the transporter and stabilizes it in an outward-open conformation. The complementarity determining regions of the antibody enter and block solvent access to the interior of the vestibule, thereby inhibiting alternating-access. NorC specifically interacts with an organic cation, tetraphenylphosphonium, although it does not demonstrate an ability to transport it. The interaction is compromised in the presence of NorC-antibody complex, consequently establishing a strategy to detect and block NorC and related efflux pumps through the use of single- domain camelid antibodies.

## Introduction

Antimicrobial resistance in superbugs acquired through multi-drug efflux facilitates the survival of pathogens against antibacterial compounds, either through direct efflux or through enhanced fitness or persistence(Blair et al., 2015; Piddock, 2006). The Gram positive superbug, methicillin resistant *Staphylococcus aureus* (MRSA) is known to cause mild skin and soft-tissue infections to severe infections like endocarditis, bacteremia, sepsis, and pneumonia(Tong et al., 2015). In the presence of penicillin, staphylococcal cultures are known to contain sub-populations of persisters that can survive the toxic effect of the antibiotic(Bigger, 1944). A major factor that aids in persistence is the enhanced activity of efflux pumps(Pu et al., 2016). Among multidrug efflux pumps, the major facilitator superfamily (MFS) constitutes an extensive array of efflux transporters with a promiscuous ability to transport a diverse array of substrates in both Gram positive and negative pathogens(Kumar et al., 2016; Reddy et al., 2012). Drug-resistant strains of *S. aureus* employ a diverse set of chromosomal and plasmid encoded MFS transporters to gain antibiotic resistance(Costa et al., 2013). Transporters like NorA, NorB, and NorC are chromosomally encoded and protect *S. aureus* against fluoroquinolones(Costa et al., 2013; Truong-Bolduc et al., 2005; Truong-Bolduc et al., 2006). QacA and QacB are plasmid encoded and provide resistance to monovalent and divalent quaternary ammonium compounds(Chitsaz and Brown, 2017; Paulsen et al., 1996a).

MFS transporters involved in multi-drug efflux primarily act by coupling efflux to proton gradients across the bacterial membrane(Paulsen et al., 1996b). MFS transporters are multi-pass integral membrane proteins comprising 12 or 14 transmembrane (TM) helices and includes drug:H^+^ antiporters (DHA) family classified as DHA1 and DHA2 depending on the presence of 12 or 14 TM helices, respectively(Reddy et al., 2012). Multiple structures of DHA1 members including MdfA, LmrP and EmrD have been solved in different conformational states that facilitate an understanding of the alternating-access in DHA members through the rocker-switch mechanism(Debruycker et al., 2020; Nagarathinam et al., 2018; Yin et al., 2006). However, there is no representative structure for the DHA2 members that comprise well-studied transporters including QacA/B, Tet38, NorB & NorC. Besides *S. aureus*, numerous pathogens express DHA2 members to effectively counter antibacterial stress. Inhibitors that block antimicrobial efflux could enhance the efficacy of existing antibiotics, thereby serving as antibiotic adjuvants (Stavri et al., 2007) (Wright, 2016).

In this study, we report the X-ray structure of NorC, a 14 TM putative efflux transporter that represents a unique subset among MFS transporters, in complex with a Zn^2+^-bound single-domain Indian camelid antibody (ICab). The structure of the ICab-NorC complex was solved at a resolution of 3.6 Å. Originally isolated as a crystallization chaperone for NorC, the ICab interacts with NorC through the insertion of CDR loops into the vestibule to effectively lock the transporter in an outward-open state. The transporter does not display a direct ability to transport fluoroquinolone or monovalent cations but has an ability to specifically interact with tetraphenylphosphonium (TPP), a monovalent cationic antibacterial whose interactions are blocked by the presence of the ICab. We anticipate that the structure of the DHA2 member NorC would facilitate the investigation of other DHA2 members in diverse pathogens and the identification of single-domain antibodies that block efflux can be explored as a novel paradigm for efflux pump detection and inhibition.

## Results and Discussion

### NorC represents a unique subset among DHA2 members

Phylogenetic analysis reveals that NorC and related sequences form a separate clade among MFS transporters implicated in multi-drug efflux (Figure 1A). Despite a clear prediction of 14 TM helices, NorC/NorB-like transporters differ substantially in comparison to the DHA2 members of the QacA-like transporters and the 14 TM proton-coupled oligopeptide symporters (POTs). Among the members of this subset, a high degree of sequence conservation is observed (∼58-88%) (Supplemental Figure 1S-1). Given the ∼70% sequence identity of NorC with NorB, it is highly likely that they would be performing very similar roles to aid in antimicrobial resistance. It is observed that NorB can aid in the survival of *S. aureus* in an abscess environment and is also overexpressed in persister populations of antibiotic resistant *S. aureus* strains (Dawan et al., 2020; Ding et al., 2008; Truong-Bolduc et al., 2006). Interestingly, we discovered that NorC/NorB-like transporters lack the typical protonation and conserved motif C residues that are a characteristic of drug:H^+^ antiporters (Varela et al., 1995). While conventional DHA2 members retain one or more negatively charged residues for protonation driven efflux, NorB and NorC lack negative charges facing the transport vestibule. For instance, Asp34 (TM1), a conserved residue amongst all the known DHA2 members for substrate recognition and protonation, is replaced by a glutamine in the NorB/NorC clade. These changes may have significant consequences on the transport properties of NorC. Also, most members related to NorC retain an amidohydrolase in the open reading frame alongside the transporter gene, which could be involved in processing the substrates transported by NorC.

**Figure 1.**
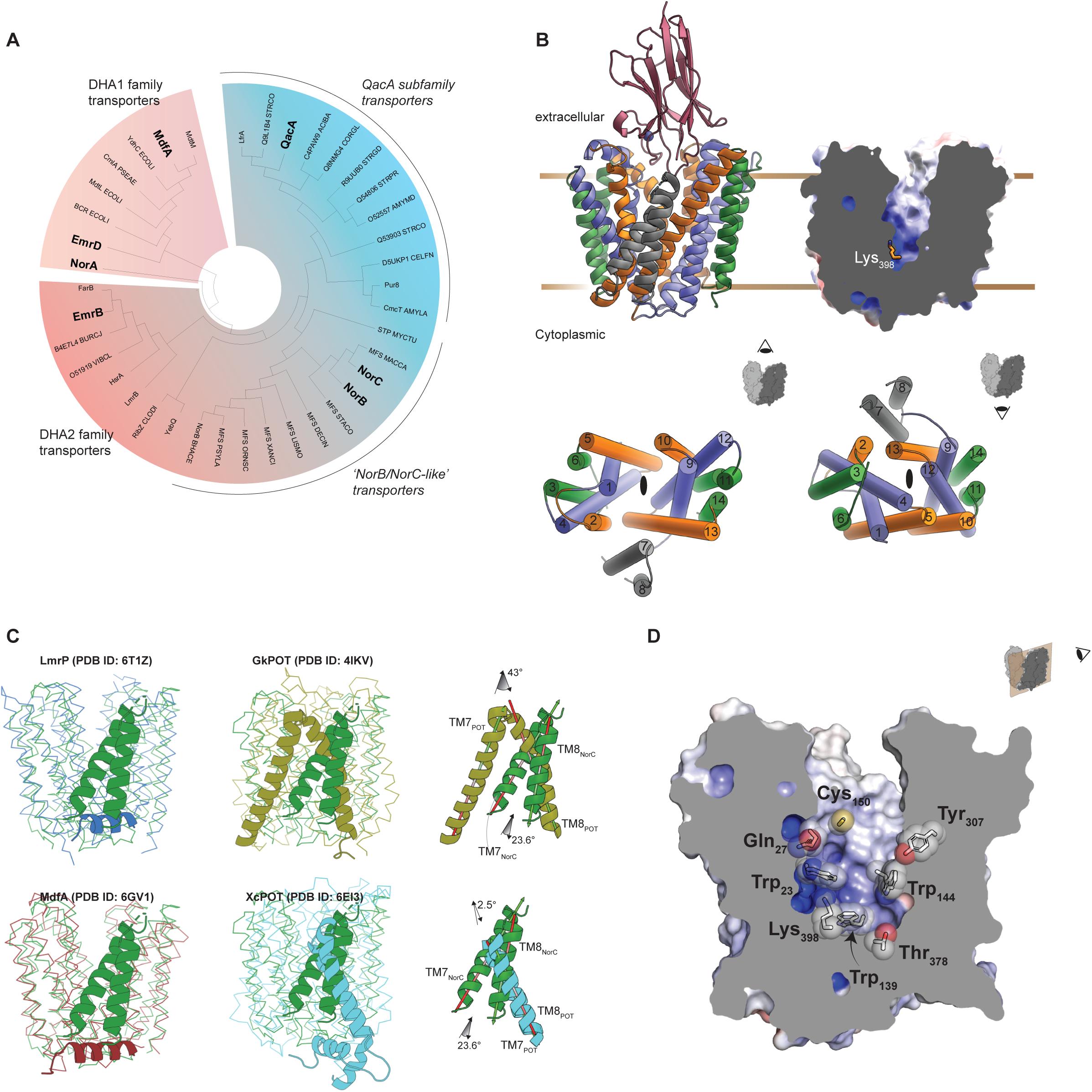
Phylogeny and X-ray structure of NorC. **A**. (top) Radial phenogram of DHA1 and DHA2 transporters. Characterized DHA2 transporters and their close homologs have been categorized with QacA/EmrB and their homologs for their sequentially coherent residues. **B**. (top-left) Side view of NorC bound to the camelid antibody ICab in outward-open conformation. The transmembrane helices have been grouped in repeats of three to depict pseudo-two fold symmetry; vestibule lining TM helices 1, 4, 9 and 12 are in blue, 2, 5, 10 and 13 lining the domain interface in orange, 3, 6, 11 and 14 in green, and helices 7 and 8 are in grey. (top right) APBS electrostatic representation showing positively charged region of the vestibule near the cytoplasmic end of NorC, owing to K398 (shown as sticks). (bottom) Note that TM7 and TM8 lie outside the primary helical bundles that surround the vestibule. **C**. Structural superposition of LmrP (blue, RMSD = 3.0 Å), MdfA (brown, RMSD = 2.5 Å), GkPOT (olive, RMSD = 4.4 Å) and XcPOT (cyan, RMSD = 4.2 Å) with NorC; angular differences between TMs 7 and 8 of NorC and those of POTs are shown in the same superposition. **D**. Solid Section of NorC showing vestibular environment.

### Crystallization and structure determination

The WT (wild-type) NorC protein was heterologously expressed and purified from *E. coli* membranes and crystallized in complex with a Zn^2+^-bound ICab that was identified and isolated in an earlier study(Kumar et al., 2020) (Supplemental Figure 2S-1). Despite obtaining single crystals, only a minor fraction would diffract and were susceptible to radiation damage. Multiple datasets were merged and scaled together to obtain a complete dataset to a resolution of 3.7 Å. The phases were estimated through Se-SAD phasing (described in methods). The resulting electron density was subjected to density modification, followed by manual model building and refinement. All the 14 TM helices could be modeled, and the main chain was traced with the help of selenium peaks resulting in the correct assignment of amino acids into the density. Multiple rounds of model building and refinement iteratively led to a significantly improved electron density map (Supplemental Figure 2S-2). A mutation in TM13, K398A allowed a marginal improvement of data quality to 3.6 Å with improved side-chain densities in the molecule leading to the refinement of the structure to acceptable R-factors (Table 1).

**Table 1.** Crystallographic Data and Refinement Statistics.

| ***Data Collection Statistics*** | **WT NorC** | **NorC K398A** |
| --- | --- | --- |
| X-ray source | APS NE-CAT 24ID-C | APS NE-CAT 24ID-C |
| Temperature | 100 K | 100 K |
| Datasets merged * | 8 | 6 |
| Space group | $P2_1$ | $P2_1$ |
| Cell dimensions<br>$a, b, c$ (Å)<br>$\alpha, \beta, \gamma$ (°) | 71.21, 139.87, 118.03<br>90, 106.03, 90 | 70.07, 139.14, 115.55<br>90, 105.7, 90 |
| Wavelength (Å) | 0.97910 | 0.97918 |
| Resolution (Å) | 49.18-3.65 (3.90-3.65) | 139.45-3.60 (3.85-3.60) |
| Total number of observations | 431001 (68355) | 426838 (45048) |
| Unique Reflections | 24592 (4248) | 24925 (4460) |
| Multiplicity | 17.5 (16.1) | 17.1 (10.1) |
| Data Completeness | 99.1 (95.2) | 99.9 (99.3) |
| Mean I/σI | 8.0 (1.0) | 8.5 (1.1) |
| R<sub>pim</sub> (%) | 4.8 (105.7) | 4.3 (87.9) |
| CC<sub>1/2</sub> | 0.997 (0.563) | 0.998 (0.486) |
| Anomalous completeness (%) | 99.1 (95.2) |  |
| Anomalous multiplicity | 8.8 (8) |  |
| DelAnomalous CC<sub>1/2</sub> | 0.476 (0.022) |  |
| ***Refinement Statistics*** |  |  |
| Resolution (Å) | 48.96 (3.65) | 111.68 (3.60) |
| Unique reflections | 23409 | 23566 |
| R<sub>work</sub> (%) / R<sub>free</sub> (%) | 29.8/31.4 | 31.1/31.8 |
| Non-hydrogen<br>atoms/molecules |  |  |
| Protein | 7567 | 7495 |
| Zn | 2 | 2 |
| Average B-factor (Å<sup>2</sup>) |  |  |
| Protein | 160.50 | 174.72 |
| Zn | 112.60 | 118.12 |
| RMS bond lengths (Å) | 0.0143 | 0.015 |
| RMS bond angles (°) | 1.84 | 1.79 |
| Ramachandran plot |  |  |
| Favored (%) | 91.61 | 90.97 |
| Allowed (%) | 8.39 | 9.03 |
| Outliers (%) | 0.00 | 0.00 |

\*Data were obtained by scaling together multiple datasets that were collected on different crystals. The highest-resolution shell used in the final refinement is shown in parentheses.

### X-ray structure of NorC bound to single-domain camelid antibody

The structure of NorC retains a typical MFS fold and all the 14 TM helices of NorC could be modeled unambiguously in the electron density, although loop regions connecting TM6 with TM7 (189-198), TM7 with TM8 (223-226) and TM8 with TM9 (247-257) were missing due to inherent disorder (Supplemental Figure 2S-2). The TM helices are organized in two six-helix bundles, a conserved feature of MFS transporters, with the linker between the helical domains forming two additional TM helices leading to a 6+2+6 arrangement (Figure 1B). The helices 1 to 6 and 9 to 14 are related by a pseudo-two fold symmetry that facilitates conformational transitions through rocker-switch mechanism (Drew and Boudker, 2016). The arrangement of the six-helix bundles is consistent with the organization in MdfA and LmrP structures in the outward-open state (Debruycker et al., 2020; Nagarathinam et al., 2018). The MFS members, proton-coupled oligopeptide transporters (POTs) that are involved in symport of di or tri-peptides, also comprise 14 TM helices (Doki et al., 2013; Solcan et al., 2012). However, all the POT structures known thus far were elucidated in the inward-open conformation unlike NorC, which is in the outward-open conformation. The relative positions of the additional TM helices 7 and 8 differ substantially in their orientation in NorC, in comparison to equivalent helices in POTs (Figure 1C). Despite these differences, the presence of two additional helices as an insertion in the linker connecting the six-helix bundles among distantly related NorC and POTs makes this a consistent organization among 14TM MFS members. The TMs 7 and 8, in NorC, are observed to occlude a wide opening between the helical bundles towards the outer leaflet of the membrane lined by TMs2 and 13 (Supplemental Figure 2S-3A). When compared to the Hoechst bound LmrP structure(Debruycker et al., 2020), the position of TMs7 and 8 clashes with Hoechst, thereby indicating the ability of TMs 7 and 8 to regulate lateral entry and binding of substrates in the vestibule (Supplemental Figure 2S-3B).

The structure of NorC is in an outward-open conformation with entry into the vestibule blocked by the ICab that was used as a crystallization chaperone (Figure 2A). Both aromatic and polar residues line the vestibule creating hydrophobic pockets as well as regions that can allow interactions with polar or charged substrates (Figure 1D). Unlike QacA, NorC lacks any negatively charged residues within the vestibule although a single cationic charge at Lys398, is observed in the cytosolic half of TM13 whose side chain is positioned facing the vestibule (Figure 1B). Incidentally, lysines can also couple protonation-driven transport as observed in Na^+^/H^+^ antiporters and aminoacid, polyamine and organocation (ApcT) transporters(Shaffer et al., 2009; Uzdavinys et al., 2017). The location of K398 coincides with the position of E407 in TM13 of QacA homology model where it was observed to be vital as both a general protonation site and substrate recognition site for certain substrates(Majumder et al., 2019). The vestibule also has a substitution of Q27 instead of acidic residues (D34) commonly observed in DHA members that transport cations including QacA and MdfA.

**Figure 2.**
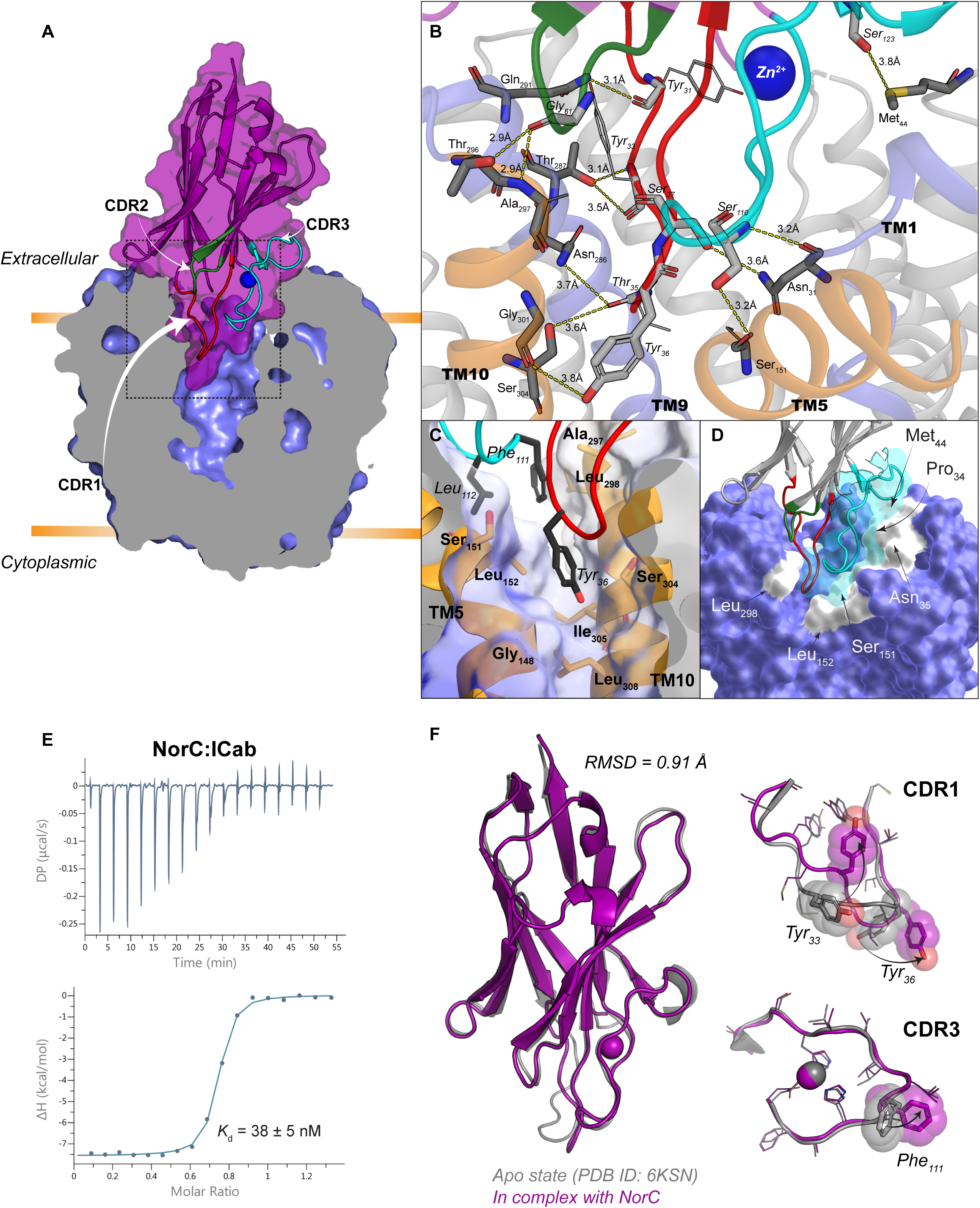
ICab interacts with NorC in an outward-open state. **A**. Solid section of NorC (light blue) in complex with ICab (purple). CDRs 1-3 are shown in red, green and cyan loops respectively, with Zn^2+^ as blue sphere. ROI in dashed border is magnified in **B**. with interchain H-bonds (dashed lines, lengths labelled) and participating residues (in sticks, light grey for ICab and dark grey for NorC; in lines shown for those not participating via their side chains, for clarity). NorC helices interacting with ICab through H-bonds are shown as cartoon. **C**. Tyr36 (CDR1) and Phe111 and Leu112 (CDR3) involved in extensive hydrophobic interactions at the interface of the N- and C-terminal domain of NorC are shown with participating residues (sticks) and complementary NorC contour (transparent surface). **D**. CDR3 (cyan cartoon and surface) forming a near perfect shape complementary, with interacting residues from NorC (white surface patches) involved in polar and VdW interactions. **E**. ITC profile (differential power, top panel; binding isotherms with integrated peaks normalized to moles of injectant and offset corrected, bottom panel.) of NorC titrated with ICab. *K*_d_ = 38 ± 5 nM. **F**. Structural alignment of ICab in native state (grey) and when bound to NorC (purple). CDR1 and CDR3 display the conformational changes that residues (sticks and transparent spheres) undergo during complexation (arrows).

Most MFS transporters are characterized by the presence of conserved motifs that are characteristic of MFS members despite weak sequence identities. In NorC, the motif A with a consensus sequence G_66_X_3_D_70_(K/R)XGR_74_X(K/R) is well-conserved and consistent with other MFS transporters (Supplemental Figure 1S-1). The Asp70 is involved in a salt bridge interaction with Arg74 of the same motif (Supplemental Figure 2S-4). Arg105 in TM4 forms the motif B and interacts with the main chain carbonyl groups of A26 (TM1) and occupies a similar position in comparison to its equivalent residue Arg112 in MdfA. The TM5 helix comprises motif C, which is characteristically present in antiporters and includes a stretch of glycine residues interspersed with a GP dipeptide (GX_8_GX_3_GPX_2_GG) (Varela et al., 1995). This motif is consistently present across numerous antiporter sequences and is absent in the case of symporters and uniporters(Varela et al., 1995). Incidentally, NorC retains a bulk of the glycine residues although the GP dipeptide is substituted by -CS-dipeptide that can have profound implications on its transport characteristics.

### ICab interacts with NorC in the outward-open state

The vestibule of NorC is open to the extracellular side although solvent access is limited through the interactions with ICab (Figure 2A). Nanobodies derived from llamas have proven to be powerful tools in studying integral membrane protein structures(Manglik et al., 2017). A few transporters like LacY have been crystallized in complex with nanobodies that bind to the extracellular face of the transporter. However, the nanobody bound LacY retains sugar interactions in the binding pocket(Jiang et al., 2016; Kumar et al., 2018a). Interestingly, the ICab has a longer CDR1 loop compared to other camelid and llama antibodies and the CDR3 loop harbors a unique Zn^2+^- binding site. Disturbing the Zn^2+^-binding site led to a loss of interactions to NorC suggesting that it is vital to retain interactions with NorC(Kumar et al., 2020). The CDR1 loop has an additional -STYS-motif that forms a β-turn and inserts deep into the vestibule of NorC, to nearly half its depth (Figure 2A) forming multiple polar and hydrophobic interactions with NorC (Figure 2B). The motif wedges between symmetry-related helices TM5 from the N-terminal domain and TM 9 and TM 10 from the C-terminal domain. TM5 of MdfA undergoes angular shifts of 15° and a clockwise twist of 45° while changing from inward-open to outward-open conformation to facilitate alternating-access(Nagarathinam et al., 2018). The TM5 of NorC is blocked by the presence of ICab whose CDR1 as well as CDR3 interacts with it. The presence of ICab CDR1 loop in the NorC vestibule further prevents the TM9 from curving as observed in the equivalent helix in MdfA(TM7) and restrains it as a linear helix (Supplemental Figure 2S-5). The CDR3 also interacts closely with residues in TMs 1, 2 and 5 towards the extracellular face (Figure 2D), which undergo substantial shifts during the rocker-switch motion. While the epitope residues do not display any interaction with the Zn^2+^ ion directly, it is evident from the NorC-ICab complex that Zn^2+^-binding allows the CDR3 to have a conformation that facilitates high-affinity interactions with NorC. The residues Phe111 and Leu112 of ICab also wedge in the gap between TMs 5 and 10 (Figure 2C).

The ICab used in this study displays a high affinity of 38 nM (Figure 2E) and interacts specifically to NorC without any cross-reactivity to NorB, despite their close similarity, as observed in FSEC experiments (Figure 3A, B). A comparison of the ICab crystal structures in the NorC-bound and free forms reveals subtle conformational changes in the CDR3 loops when it interacts with the antigen. The β-turn undergoes a straightening in its position and the Thr35 undergoes a 3.0 Å shift in its Cα position. Similarly, the Tyr36 in the β-turn undergoes a displacement of its phenol side chain by 3.3 Å to facilitate its wedging between TMs 5 and 10 (Figure 2F). The CDR3 largely retains a conformation in the bound state similar to that of the free ICab with a minor displacement of about 1.5 Å in the region between 109 and 115. However, the side chain of Phe111 undergoes a massive shift in the χ1 torsion angle with 148° rotation to interact with NorC residues Leu152 and Leu298.

**Figure 3.**
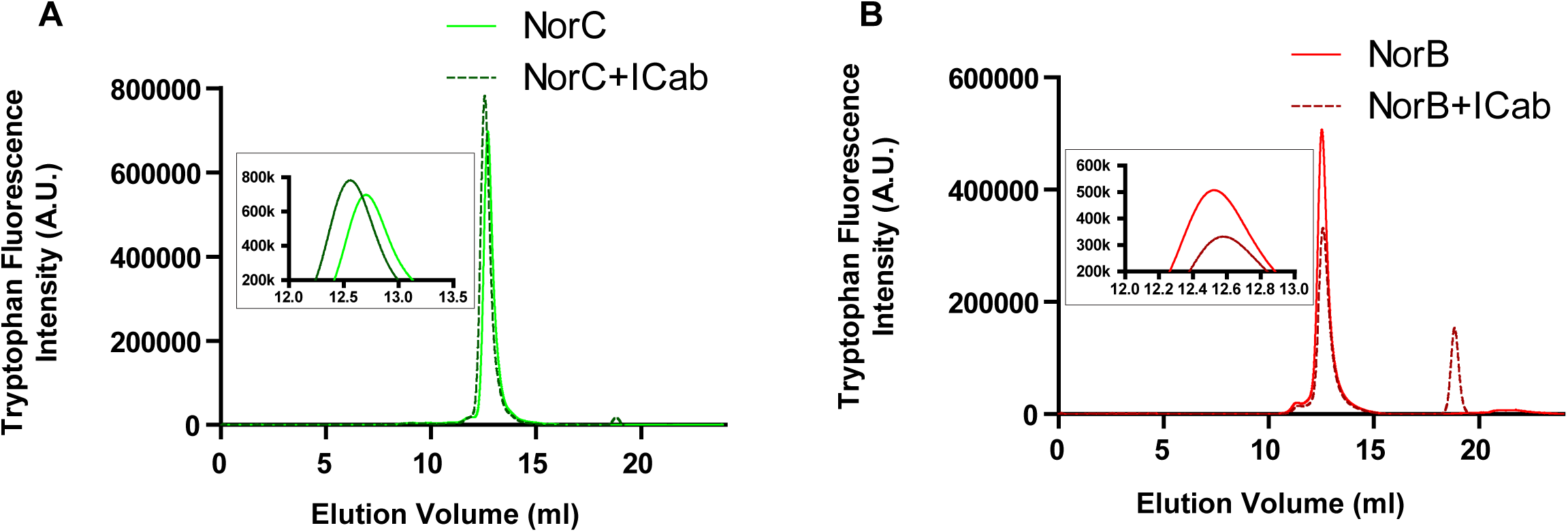
ICab sepcifically interacts with NorC. **A**. Tryptophan fluorescence trace of NorC in FSEC shifts to a slightly higher molecular mass when complexed with ICab. **B**. NorB does not display a shift indicating a lack of interaction with ICab.

### ICab blocks access to antibacterial compounds

The ability of ICab to interact with the vestibule results in a massive reduction of solvent accessibility within the vestibule of NorC, likely compromising substrate interactions (Figure 4A). The natural substrates of NorB/NorC-like transporters are unknown. As NorC is reported to be involved in efflux of fluoroquinolones, we performed survival assay of *E. coli* expressing NorC in presence of fluoroquinolones. To our surprise, we did not observe a direct efflux of fluoroquinolones with NorC (Supplemental Figure 1S-2). While screening for potential substrates that can interact with NorC, we discovered specific interactions between NorC and TPP with a *K*_d_ value of 4 μM (Figure 4B). Although TPP interactions with both NorB and NorC K398A led to a nominal enhancement of stability (Δ*T*_*m*_ = +1.5°C and +2.9°C respectively) (Supplemental Figure 4S-1), WT NorC lacks the ability to transport TPP (Supplemental Figure 1S-2). This is plausible in the context of other MFS transporter structures like LmrP, which has a detergent molecule, a phospholipid and a cationic dye (Hoechst) bound in distinct subsites within the vestibule in LmrP(Debruycker et al., 2020). We suggest that TPP, given its multiple phenyl groups could stack and interact with the aromatic side chains within the NorC vestibule. The interactions of TPP with NorC are clearly blocked in the presence of ICab causing a complete loss of interactions of the antibacterial compound with NorC (Figure 4B). In doing so, the ICab-NorC complex, resembles a ‘bottle-cork’ that prevents substrate interactions and enforces a lock on the conformational changes within the transporter.

**Figure 4.**
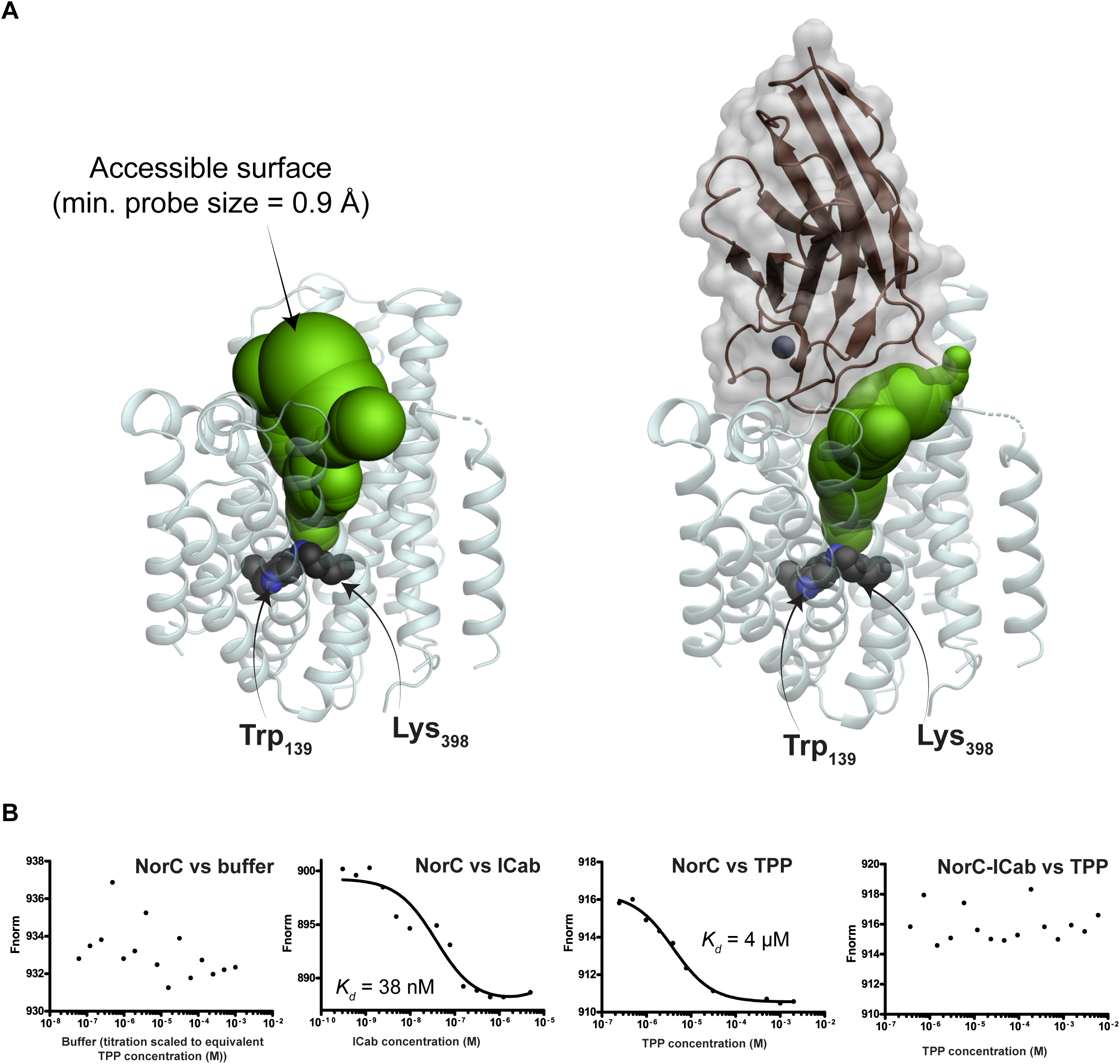
ICab interaction alters solvent accessibility of NorC vestibule. **A**. Water accessible surface (green spheres) in NorC’s vestibule starting from Trp139/Lys398 (black spheres), in the absence (left) and presence (right) of ICab. The outward open conformation completely blocks accessibility from the cytosolic side of NorC. **B**. Microscale thermophoresis profiles for NorC and tetraphenylphosphonium in the presence and absence of ICab. *K*_d_ = 4 ± 0.8 μM for NorC and TPP.

## Conclusions

NorC structure is the first representative structure of the DHA2 family and reveals the architecture of this distinct class of transporters within the MFS fold. While the true substrate of NorC is not known, both NorC and its closely related homologue, NorB are observed to protect the superbug against fluoroquinolones, which are broad-spectrum antibiotics. The lack of direct transport of fluoroquinolones through NorC suggests that the transporter could be aiding in reducing antibiotic stress through alternate mechanisms including improved fitness and persistence, as observed in the case of NorB (Ding et al., 2008) (Dawan et al., 2020). The ability to detect and block efflux transporters is proposed as a promising strategy to aid and improve the efficacy of existing antibiotics. The ICab characterized in this study provides a high fidelity detection tool to observe the presence of NorC in *S. aureus* populations. Its ability to wedge deeply in the NorC vestibule and lock it in an outward-open conformation serves as a proof-of-principle to use ICabs or nanobodies as efflux pump inhibitors and a potential strategy to overcome antimicrobial resistance.

## Acknowledgements

The authors would like to thank Dr. Rakesh Ranjan, NRCC and Dr. M. Ithayaraja for help with NorC immunization and ICab screening through yeast surface display, respectively. The authors would like to thank Prof. B. Gopal for sharing the genomic DNA of *S. aureus* strains and access to PEAQ-ITC. The authors would like to thank the staff of northeastern collaborative access team (NECAT), Advanced Photon Source, particularly Dr. Surajit Banerjee for access and help with data collection. We thank the beamline staff at the Elettra XRD2 particularly Dr. Babu Manjashetty and Dr. Annie Heroux for beamline support. Access to the XRD2 beamline at the Elettra synchrotron, Trieste was made possible through grant-in-aid from the Department of Science and Technology, India, vide grant number DSTO-1668. We would like to thank the staff of PX beamline of the Swiss Light Source for access and support.

## Author contributions

SK and AA purified NorC and ICab. SK and AA optimized the crystallization of NorC-Icab complex and selmet derivatization of NorC. SK solved and built the structure of NorC with inputs from AP. AA, SK performed the microbial growth assays and binding assays with TPP, respectively. AG and RMN cloned, isolated and optimized the behavior of NorC. AP designed the project. SK and AP wrote the manuscript with inputs from all authors.

## Data availability

The atomic coordinates and structure factors of WT NorC and NorC-K398A are deposited in the Protein Data Bank with PDB IDs 7D5P and 7D5Q, respectively.

Correspondence and requests for materials should be addressed to AP.

## Methods

### Sequence alignment and Phylogenetic analysis

NorC (Uniprot ID A0A0E1ACG1) was used to search for closely related homologous sequences in non-redundant sequence database. To search for more homologous sequences amongst the known DHA family members, transporter classification database was used. The alignment was carried out using clustalw program with iterative HMM clustering. The output properly aligned the TMs of all the query proteins –a benchmark we used to check the alignment accuracy for this highly divergent dataset. The alignment was used further for functional and phylogenetic analysis. The evolutionary history was inferred by using the maximum likelihood method and JTT matrix-based model(Jones et al., 1992). The tree with the highest log likelihood (−35519.34) is shown. Initial tree(s) for the heuristic search were obtained automatically by applying Neighbor-Join and BioNJ algorithms to a matrix of pairwise distances estimated using the JTT model, and then selecting the topology with superior log likelihood value. A discrete Gamma distribution was used to model evolutionary rate differences among sites (2 categories (*+G*, parameter = 3.0280)). This analysis involved 39 amino acid sequences. There were a total of 611 positions in the final dataset. Evolutionary analyses were conducted in MEGA X(Kumar et al., 2018b).

## Survival Assays

To check for phenotype (resistance against antibacterials), C41 cells (transformed with IPTG inducible *pET16b-norC* or *pET16b-norA)* and JD838 cells (transformed with L-arabinose inducible *pBAD-norC, pBAD-empty* or *pBAD-qacA*) were grown as primary culture overnight. A secondary culture was inoculated using this and split into two parts: one treated for induction of protein expression (using 0.2mM IPTG or 0.05% (w/v) L-arabinose) and the other left untreated. The induction was done when the O.D.s_600nm_ reached 0.4 A.U. The cultures were further grown till all of them had an O.D. _600nm_ between 1.5-2.0 A.U.; the cultures were then diluted accordingly to bring their O.D.s to 1.0 and spotted in 10 fold serial dilutions on 1.5% (w/v) agar plate containing 2% (w/v) luria bertani (LB) broth and 100 μg/ml Ampicillin with/without inducer in the media. Control plates containing none of the antibacterials but ampicillin were spotted similarly with/without inducer. The cells were allowed to grow overnight at 37°C. These assays were replicated independently with n=5.

### Cloning, expression and purification of NorC

Full length *norC* gene was cloned into pET-16b vector between restriction sites NcoI and KpnI with 8X-His tag at its C-terminus. The *E. coli* BL21-C41 cells were transformed with the cloned vector containing ampicillin resistance cassette. A primary culture was grown in LB-broth (HiMedia) for 10-12 hours at 37 °C. Primary culture was used to inoculate large scale culture (2-3 l) that were grown at 37 °C until the OD_600_ reached 0.4-0.5 following which protein expression was induced with 0.2 mM Isopropyl-β-D-1-thiogalactopyranoside (IPTG). The cells were further grown for 12-15 hrs at 20°C. Cells were harvested by centrifugation and flash frozen in liquid N_2_ for storage or processed immediately. The cell pellet was dissolved in HBS (20 mM Hepes pH 7.0, and 200 mM NaCl). The cell suspension was lysed at high pressure of about 800 bar using a homogenizer (GEA Niro Soavi PandaPlus 1000 Homogenizer) and spun at ∼100000*g* for 1 hour at 4 °C. The pelleted membranes were resuspended in HBS and 1mM PMSF using a rotor-stator homogenizer, following which DDM (n-Dodecyl-β-D-maltopyranoside, Anatrace) was added to a final concentration of 20mM. The solution was nutated for 2 hours at 4°C to extract NorC into micelles, followed by ultracentrifugation at 100000 *g* for 1 hour at to remove the insoluble debris. The supernatant was incubated with Ni-NTA beads pre-equilibrated with buffer A (1mM DDM in HBS) at 4°C for 1 hour. The solution containing beads were transferred into gravity columns (Bio-rad) and washed with 50 column volumes of buffer A having 30mM imidazole. The protein was eluted in buffer A containing 300 mM imidazole and concentrated using a 30 kDa cut-off centrifugal filter (Amicon, Merck Millipore). It was further purified by size exclusion chromatography using Superdex S-200 increase 10/300 GL column (GE Healthcare) in HBS containing 4 mM n-Decyl-β-D-maltopyranoside (DM, Anatrace). The purity of the protein was analyzed by SDS-PAGE. The mutant NorC K398A protein was purified in a similar manner.

WT NorC was labelled with L-selenomethionine by growing *E. coli* BL21-C41 cells, transformed with NorC-containing vector, in M9 medium supplemented with 50 mg/L of L-selenomethionine. The purification of SeMet-labeled WT NorC was similar to that of the native protein.

### ICab generation and purification

Generation and purification of ICab was carried out in a manner described previously(Kumar et al., 2020). The ICab isolated through yeast-display screening was subcloned into pET-22b vector. *E. coli* Rosetta cells were transformed with *vhh*-containing vector. Large scale cultures were grown in LB broth 37 °C until OD_600_ reached 0.6 and protein expression was induced by adding IPTG to a final concentration of 0.5 mM; cells were further grown at 37 °C for 5-6 hours. The cells were harvested and the protein was extracted from inclusion bodies using urea denaturation followed by refolding. The refolding was done using step-wise dialysis at 4°C. The protein was further purified by size-exclusion chromatography using Superdex S-75 10/300 GL column (GE Healthcare) in HBS. The purity and integrity of the protein was checked by SDS-PAGE and MALDI-TOF.

### Site-directed mutagenesis

To perform site-directed mutagenesis of NorC, primers were designed to amplify entire plasmid carrying *norC* gene in a single step with desired point mutation. The amplicons were treated with Dpn1 at 37°C for 2-3 hours to digest the wild-type plasmid. *E. coli* Top10 cells were transformed with the mutant amplicon and plated on LB-agar plate containing 100 µg/ml ampicillin and grown overnight at 37°C. The positive clones were confirmed by DNA sequencing. The mutant proteins were purified in a manner similar to that of the wild type protein.

### Fluorescence-detection size exclusion chromatography (FSEC)

Briefly, FSEC was done using a high-performance liquid chromatography system attached to an autosampler and a multi-wavelength fluorescence detector (Shimadzu). 10μM each of WT NorC, NorC-K398A and NorB purified in DM buffer were incubated with 1:1.2 molar ratios of ICab and shifts in their elutions (measured at λ_ex_ = 395nm, λ_em_ = 340nm) in the same DM buffer were used to qualitatively determine whether or not ICab binds to them. A superdex 200 HR column was used for analyzing elution times.

### Crystallization and structure determination

Purified WT NorC (native and SeMet-derivative) and NorCK398A (native only) proteins were concentrated to 3 mg/ml using 50 kDa cut-off centrifugal filter (Amicon, Merck Millipore) and mixed with ICab in a molar ration of 1:1.2. The crystallization was carried out by hanging drop vapor diffusion method at 20 °C where protein and crystallization conditions were mixed in a ratio of 3:2 v/v. WT NorC and NorC-K398A crystallized in a condition containing 0.1 M MES pH 6.0, 50 mM NaCl, 35.7 % PEG 600, 57.1 mM CaCl_2_, 10 mM YCl_3_ and 6.0 mM CHAPSO. The crystals appeared within 3-4 days but took about 2 weeks to attain full size. The crystals were washed in the crystallization condition and flash frozen in liquid N_2_. The X-ray diffraction data sets were collected at Advanced Photon Source (APS) Synchrotron, USA. The crystals diffracted X-rays to a resolution of about 3.6-4Å.

All the data sets were indexed and integrated with XDS (Kabsch, 2010). Eight data sets were merged and scaled for SeMet-derivative WT NorC using AIMLESS (Evans, 2006) resulting in a highest resolution of 3.65Å. The space group was determined to be P2_1_ with unit cell dimensions of a=71.2 Å, b=139.9 Å, c=118.0 Å and α=γ=90° and β= 106° (Extended Data Table 1). The Matthews coefficient suggested two NorC-ICab complexes in the asymmetric unit with a solvent content of about 70 percent. SAD phasing was carried out using CRANK2 (Skubak and Pannu, 2013) in CCP4i2 (Winn et al., 2011). For the substructure determination, the highest resolution was set at 5.0 Å as anomalous signal was very weak at higher resolution. SHELEX C and D (Schneider and Sheldrick, 2002) in CRANK2 were used for substructure determination and refinement. The resulting phases were extended to a resolution of 3.65 Å and the electron density was subjected to density modification using PARROT (Cowtan, 2010). Manual model building was carried out in the density modified map using Coot (Emsley and Cowtan, 2004). Selenium positions were used as markers to trace the main chain and place amino acids.

Multiple X-ray diffraction data sets were collected for the native NorCK398A crystals. Six best data sets were merged and scaled using AIMLESS to a resolution of 3.6 Å. The structure was determined by molecular replacement using PHASER (McCoy et al., 2007) with WT NorC structure as a model. The refinement of both the structures were carried out using REFMAC(Murshudov et al., 2011) until the refinement converged. The resulting structures were validated using MOLPROBITY(Williams et al., 2018). The data collection and refinement statistics are presented in the Extended Data Table 1.

### NorC interaction with TPP

Quantification of NorC’s interaction with TPP was carried out using microscale thermophoresis (Nanotemper)(Seidel et al., 2013). The protein was labelled with red Tris-NTA dye (Nanotemper) by mixing both in an equimolar proportion. The labelled protein was mixed with TPP such that the concentration of labelled protein and TPP were 50 nM and 4 mM respectively. A total of 16 two-fold serial dilutions of TPP were prepared keeping the protein concentration constant at 50 nM. The experiment was carried out with Monolith™ NT.115 MST premium-coated capillaries. For NorC-ICab complex titrations against TPP, 50 nM of NorC was mixed with ICab in 1:3 molar ratio and the starting concentration of TPP was kept at 4 mM. For ICab titrations against NorC, 10 nM of NorC was used with a starting concentration of 5 µM for the ICab. The curves were fitted with single-site binding model.

TPP’s propensity to interact with NorB and NorC-K398A was similarly measured using differential scanning fluorimetry (DSF) using Prometheus NT.48 instrument (NanoTemper Technologies)(Kotov et al., 2019). A temperature scan was carried out between 20 and 90 °C with a scan rate of 1 °C/min. All measurements were carried out in duplicates. The first derivative of the ratio of F350/330 was plotted against temperature, and shift in *T*_*m*_ was used to determine whether or not TPP binds to the proteins.

### Isothermal Titration Calorimetry

The binding affinity of NorC with ICab was determined by ITC using Microcal PEAQ-ITC (Malvern Panalytical). Both NorC and ICab were purified in a buffer containing 20 mM Hepes, pH 7.0, 200 mM NaCl, and 4 mM decyl-β-D-maltopyranoside detergent. The titration consisted of 18 injections of 4s duration each, with first injection of 0.4 µl and all the subsequent injections of 2 µl. The time between two consecutive injections was kept at 150 s, and the sample in the cell was stirred at 750 rpm during the entire run. The concentration of NorC in the cell was kept at 25 µM and the concentration of ICab in the syringe was kept at 250 µM. The titration was carried out at a constant temperature of 25 °C. For blank, ICab was titrated against buffer, and the heat of mixing and dilution were subtracted from the titration data against NorC. All of the data were fit using Microcal PEAQ-ITC analysis software with one set of sites model(Kumar et al., 2020).

