## Supplementary Figures for "Structural basis of inhibition of a putative drug efflux transporter NorC, through a single-domain camelid antibody"

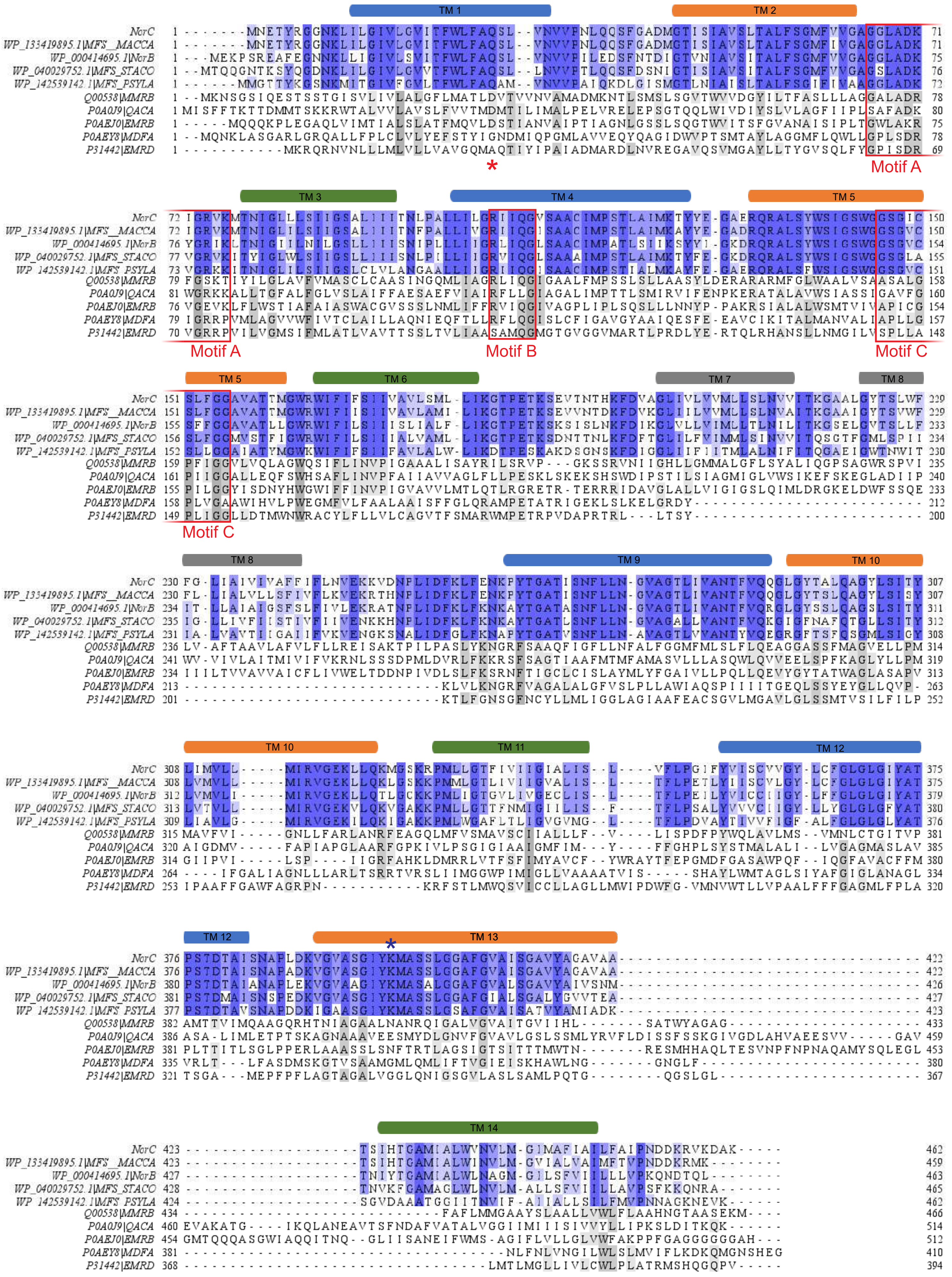

**Figure 1-figure supplement 1** | Sequence alignment of NorB/NorC-like and conventional DHA1 and DHA2 family members. Conservation is shown in blues and greys, respectively. Residue conservation declines as highlights go lighter. TM helices are identified as cylinders above the sequences. Motifs A, B and C are boxed in red. Asp34 (DHA1 and DHA2), and Lys398 (NorC/NorB subfamily) are marked with asterisks.

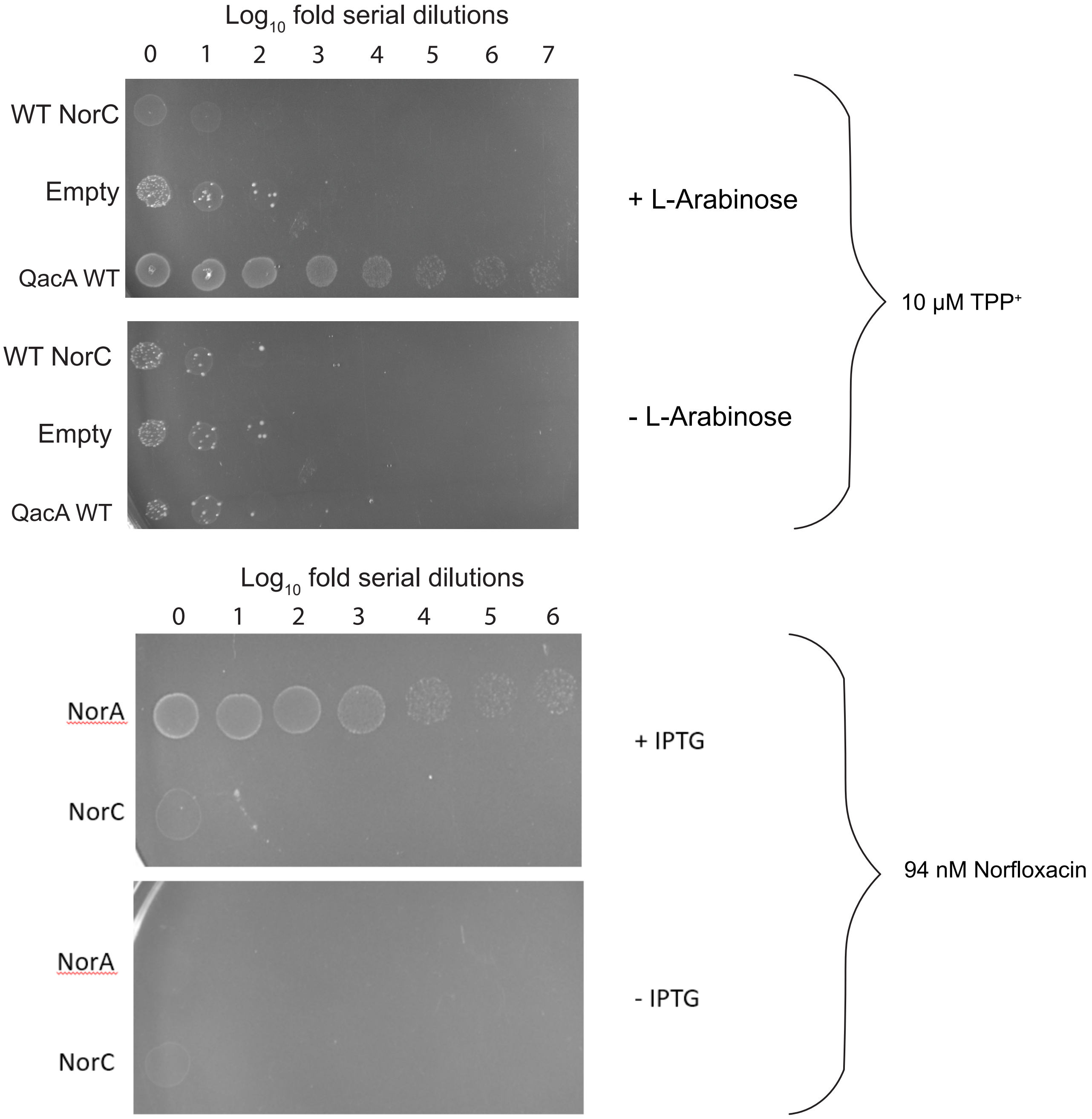


**Figure 1-figure supplement 2|** Survival assays done in the presence of tetraphenylphosphonium ($TPP^+$, using L-arabinose inducible pBAD24 expression vector) and norfloxacin (using IPTG inducible pET16b expression vector) at known $MIC_{50}$; n=5 for independent replicates.

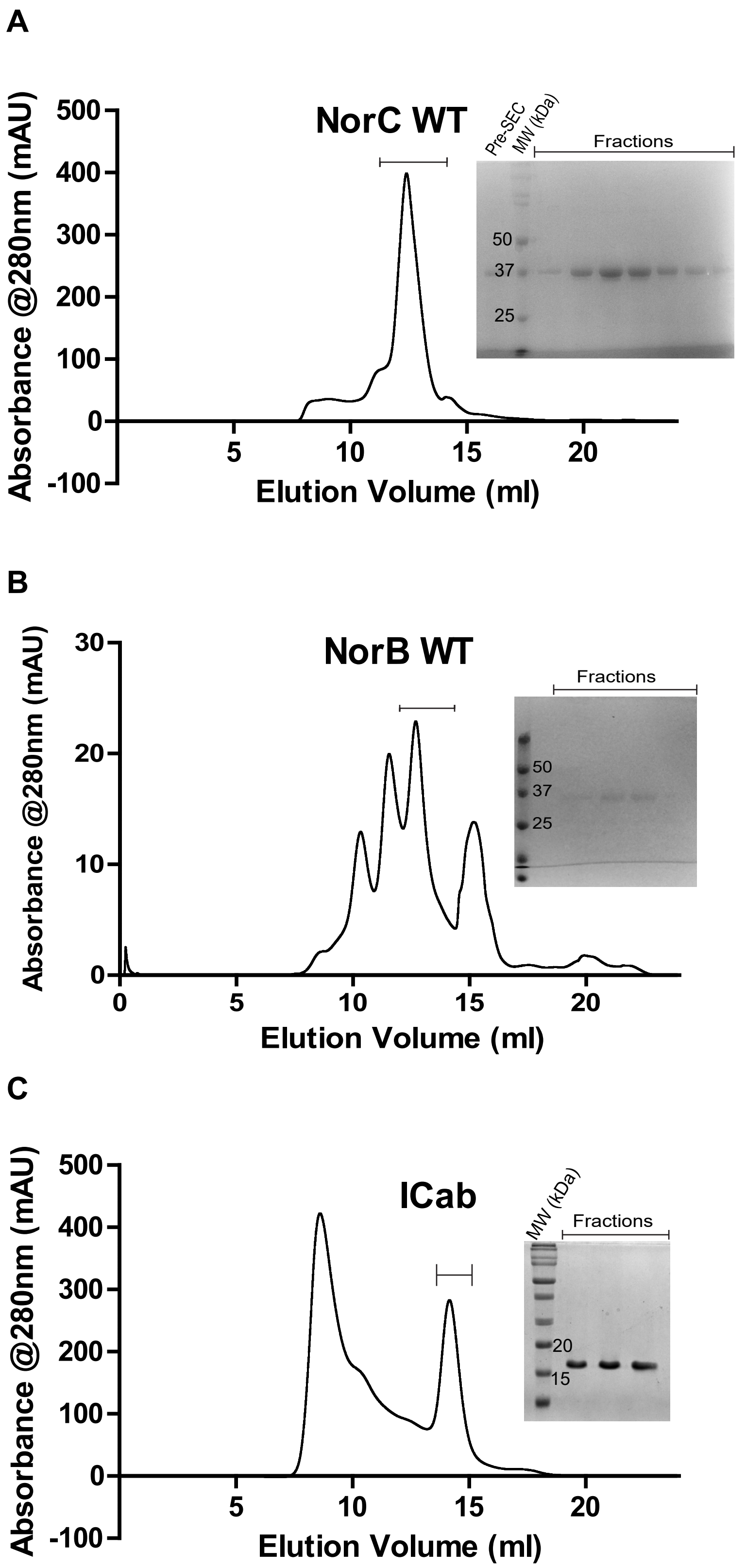


**Figure 2-figure supplement 1**| Size exclusion chromatography profiles and SDS-PAGE (insets) with corresponding fractions of **A.** NorC, **B.** NorB and **C.** ICab.

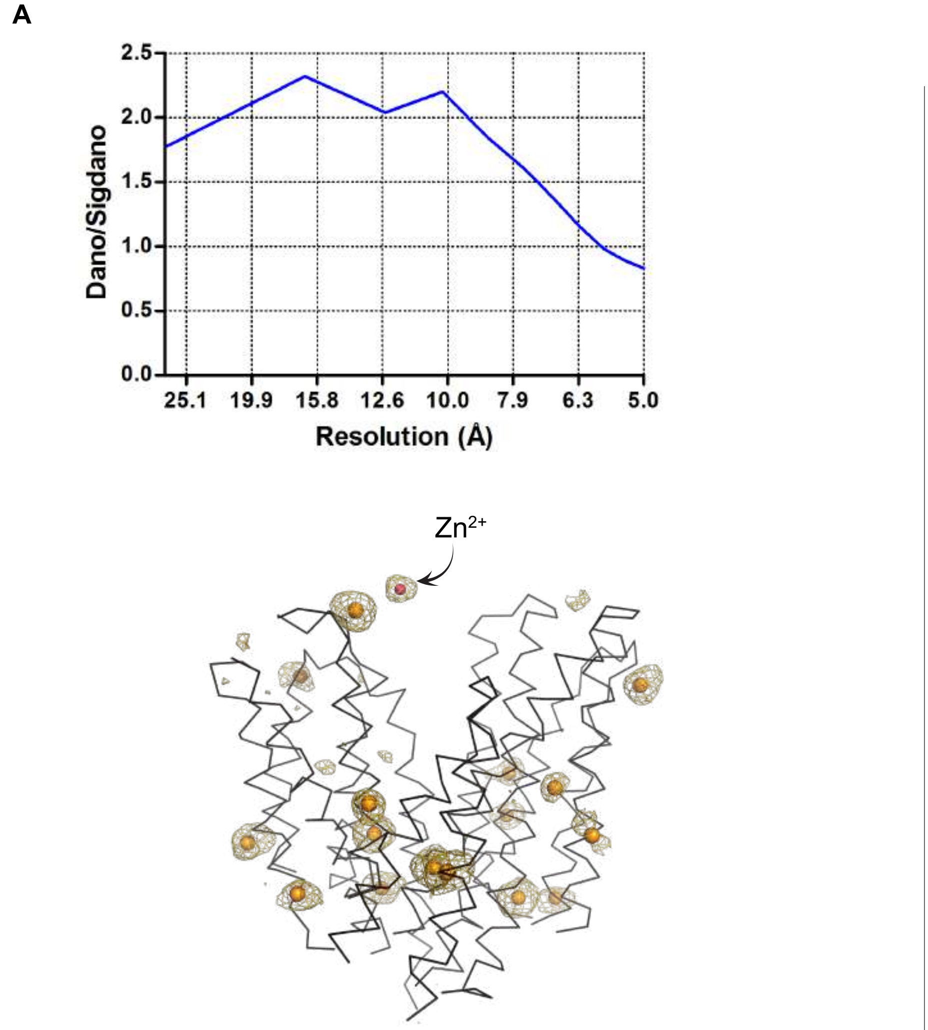


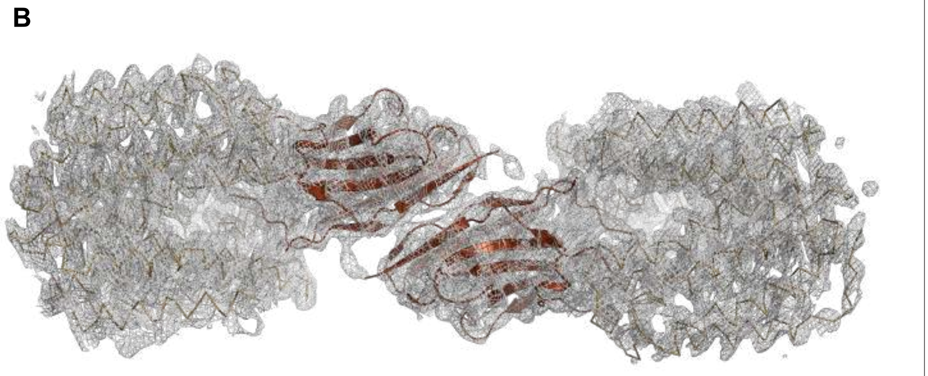


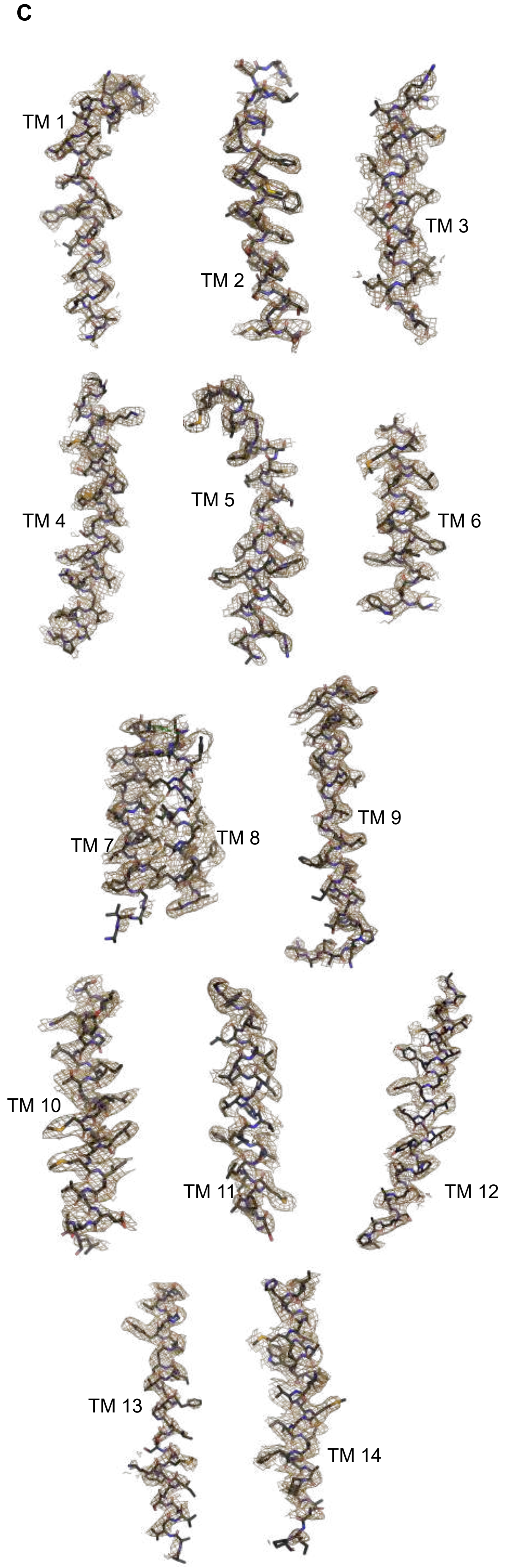


**Figure 2-figure supplement 2** | **A.** (Top) signal quality represented by $\Delta I_{Ano}/\sigma(\Delta I_{Ano})$. Anomalous phasing and substructure determination was done at 5 Å; 16 out 17 Se atom positions were determined with their anomalous densities shown in golden spheres and meshes respectively. $Zn^{2+}$ density is also seen, shown in pink sphere. **B.** ASU of NorC-ICab complex contains 2 molecules of each, shown with 2Fo-Fc map contoured at 1.0 σ; focused ROI displays interface between ICab chains (stick representation). **C.** Individual TM helices and their surrounding 2Fo-Fc map at 1.0 σ carved at 2.0 Å.

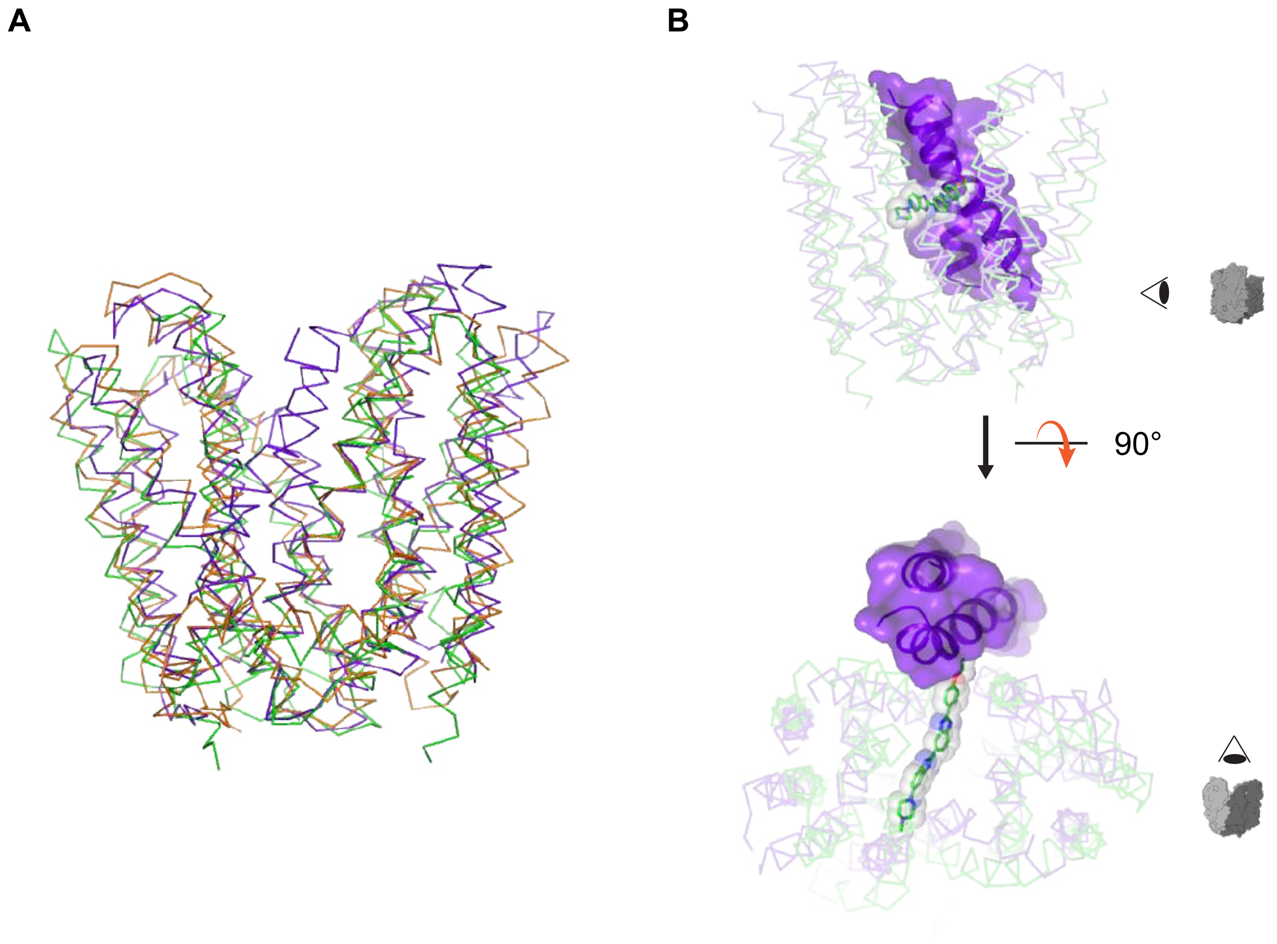


**Figure 2-figure supplement 3** | **A.** Superposition of NorC (purple), MdfA (PDBID: 6GV1, orange), and LmrP (PDBID: 6T1Z, green) structures. TMs 7 and 8 of NorC occlude a wide opening going from the vestibule to the upper half of the membrane, through a crevice made by TMs 2 and 13 **B.** Side and top views of the superposition showing the substrate Hoechst (sticks and spheres, bound to LmrP) clashing with TM7 (purple surface) of NorC.

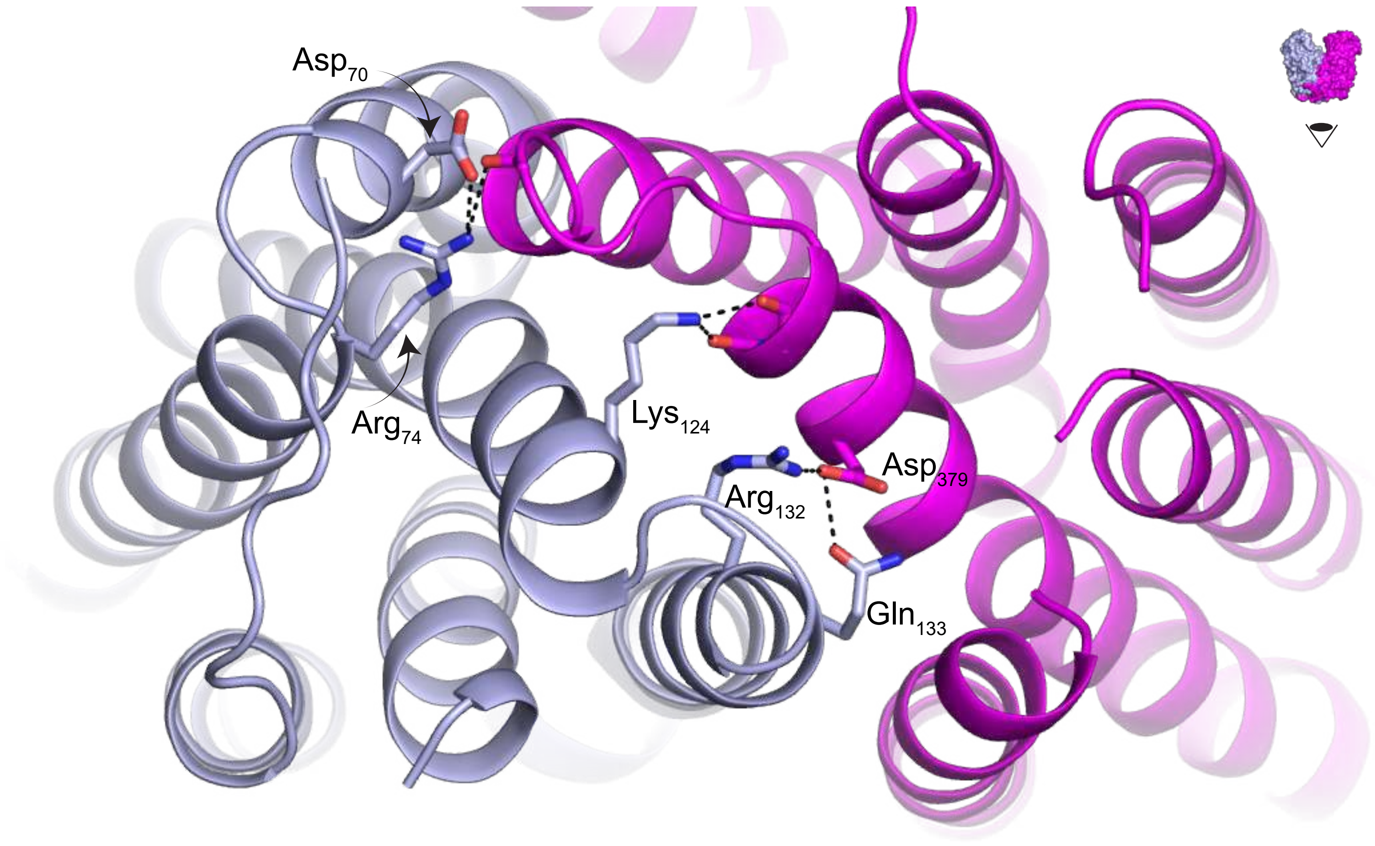


**Figure 2-figure supplement 4 |** Polar interactions between two symmetry-related halves of NorC (N-terminal in slate, C-terminal in magenta) towards the cytosolic side.

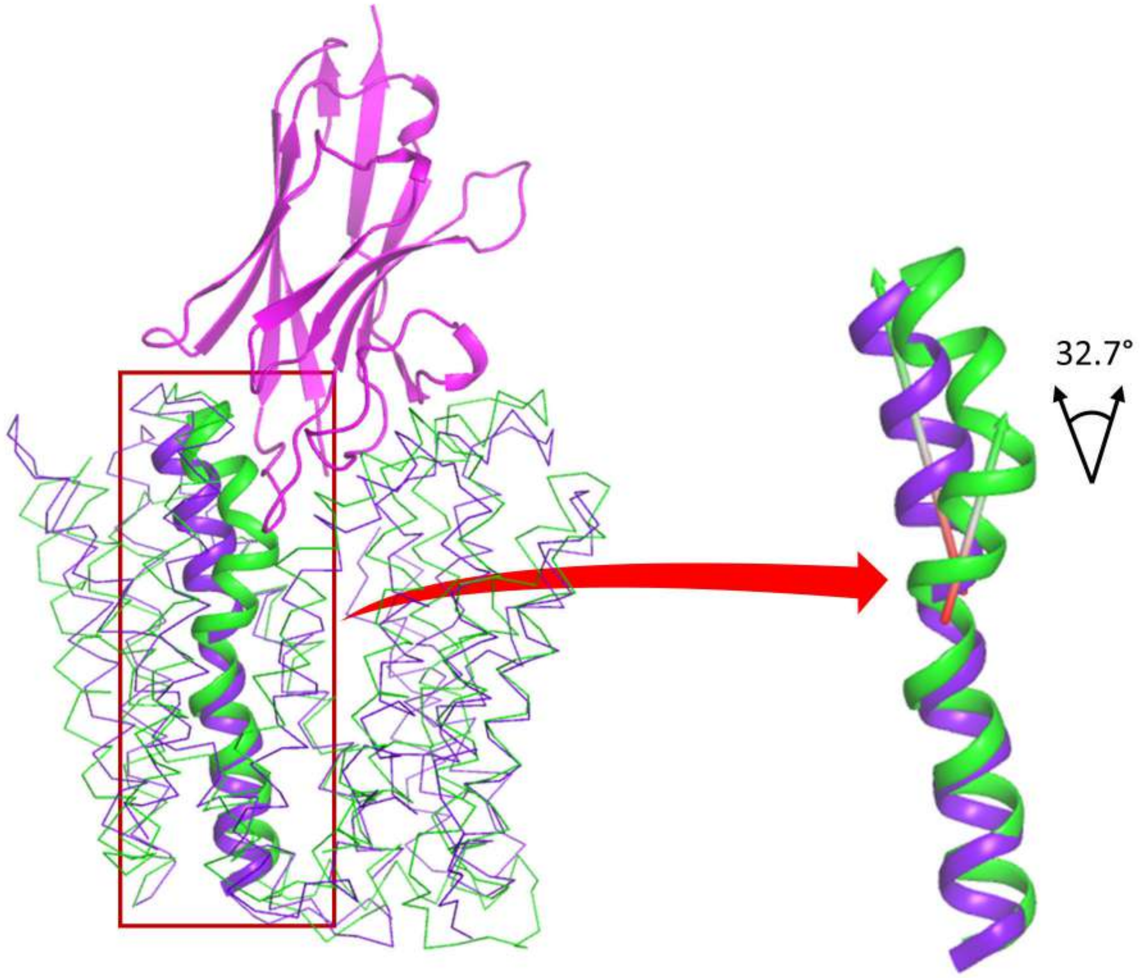


**Figure 2-figure supplement 5 |** Superposition of NorC-ICab (purple and magenta) complex and MdfA (PDB ID: 6GV1, green). TM7 of MdfA and TM9 of NorC are shown in cartoon representation; where TM9 of NorC is relatively unbent whereas TM7 of MdfA kinks to an angle of 32.7° (enlarged view, right).

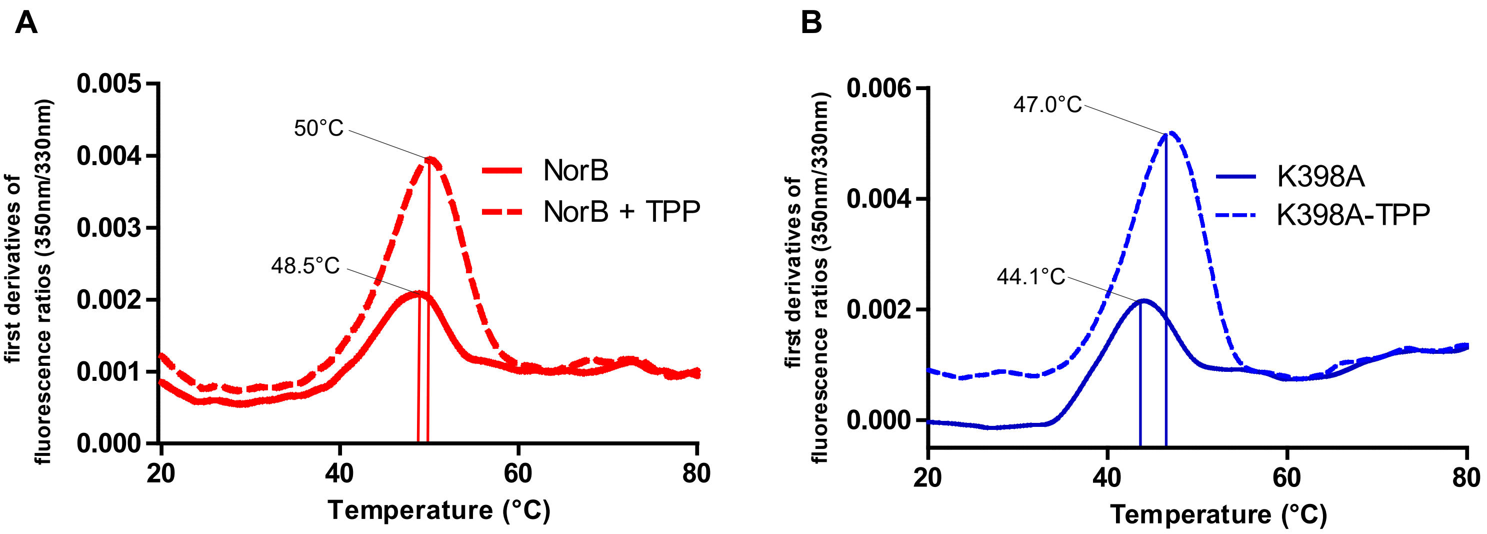


**Figure 4-figure supplement 1** | Differential scanning fluorimetric profiles of **A.** NorB in the presence and absence of $TPP^+$ ($\Delta T = 1.5°C$, n=3, technical replicates), and **B.** the same with NorC-K398A ($\Delta T = 2.9°C$, n=2, technical replicates).
